# Establishment of an *in vitro* Culture and Regeneration Protocol for the Native Grass *Polypogon australis* Brong

**DOI:** 10.1101/2025.11.12.688029

**Authors:** Javiera Venegas-Rioseco, Claudia Ortiz-Calderón, Götz Hensel, Rosanna Ginocchio

## Abstract

*Polypogon australis* (Brong.), a native Chilean grass (Poaceae), is a facultative metallophyte capable of colonizing copper mine tailing dams and adapting to saline and acidic substrates. These traits make it a promising candidate for phytoremediation of metal-contaminated soils. However, the lack of *in vitro* propagation, callus induction, and somatic embryogenesis protocols limits its use in large-scale applications and genetic improvement. This work aims to establish a reproducible *in vitro* regeneration system for *P. australis*. Mesocotyl explants were cultured on Murashige and Skoog (MS) medium supplemented with 2.5 mg L^−1^ Dicamba. Callus induction was achieved in 38.9% of explants, and 45.5% of embryogenic calli regenerated into plantlets producing leaves and radicles without requiring exogenous organogenesis-inducing hormones. The regenerated plants continued to develop further on MS + sucrose medium, confirming the totipotent capacity of mesocotyl-derived calli. The developed protocol provides a foundation for large-scale propagation and genetic transformation of *P. australis*. By overcoming propagation bottlenecks, this methodology strengthens the potential of this native metallophyte as a model for phytoremediation and future CRISPR-based biotechnological approaches to enhance copper tolerance and accumulation.

## Introduction

Native plants that colonize metal-contaminated soils and industrial solid wastes (i.e. mine tailings) have developed unique adaptations, including enhanced metal uptake, detoxification, and tolerance mechanisms, allowing them to survive and reproduce under extreme conditions [1,2]. These traits make metallophytes promising candidates for phytotechnologies such as phytoextraction, phytostabilization and phytomining, which offer eco-friendly, cost-effective, and minimally invasive strategies for soil and substrate remediation and valorization [3,4].

However, the biotechnological optimization and application of these plant-based technologies require validated and reproducible propagation protocols, such as *in vitro* technologies, that are essential for large-scale plant production, studying stress physiology, and as platforms for genetic improvement [5]. Tissue culture technology is particularly relevant for conserving and propagating native plants that are adapted to stressful environments, including those with metal-rich or saline soils [6]. Genetic engineering further expands these opportunities by enabling the enhancement of traits such as metal uptake, oxidative stress resistance, and biomass accumulation, thereby improving phytotechnologies efficiency [7]. Despite these advances, efficient *in vitro* methodologies are still lacking for many native metallophytes, limiting their broader application [8].

Among plant families, Poaceae are often prioritized for phytotechnologies due to their extensive root systems, rapid growth, and ability to accumulate metals in harvestable tissues [9]. *Polypogon australis* Brong. is a facultative metallophyte native to Chile that spontaneously colonizes abandoned copper tailing dams, accumulating Cu concentrations of 669.5 mg kg^−1^ (DW) in leaves and 223 mg kg^−1^ (DW) in roots, with a translocation factor of 3.0 [10]. Its natural adaptation to the harsh conditions of northern Chile, including saline and acidic soils, further highlights its resilience [11]. Moreover, this species tolerates additional pollutants, such as hydrocarbons [12], and maintains antioxidant enzymatic activity when exposed to copper-rich leachates [13].

Nevertheless, the absence of *in vitro* propagation, somatic embryogenesis, and regeneration protocols for *P. australis* constrains its use in phytotechnological approaches [9,14]. Tissue culture in native Poaceae is often challenged by species-specific recalcitrance and limited embryogenic responses [15]. To address these limitations, the present study establishes an efficient *in vitro* propagation system for *P. australis* using mesocotyl-derived callus induction and regeneration. This methodology lays the foundation for large-scale propagation and provides a basis for genetic transformation, ultimately enhancing the potential of this species for being used in phytotechnologies and biotechnology [7].

## Materials and Methods

### Plant material and seed sterilization

Mature seeds of *Polypogon australis* were collected from plants growing on a copper mine tailings storage facility in northern Chile (Planta Manuel Antonio Matta, Atacama, Chile; 27.3° S, 70.3° W) (Fig. 1A) [16]. Surface sterilization followed Acemi and Özen [17] with modifications. Seeds were immersed in 70% (v/v) ethanol for 1 minute, followed by 1% (w/v) sodium hypochlorite solution containing a drop of Tween™ 20 for 10 minutes, and then rinsed three times with sterile distilled water (Fig. 1B).

**Figure 1.**
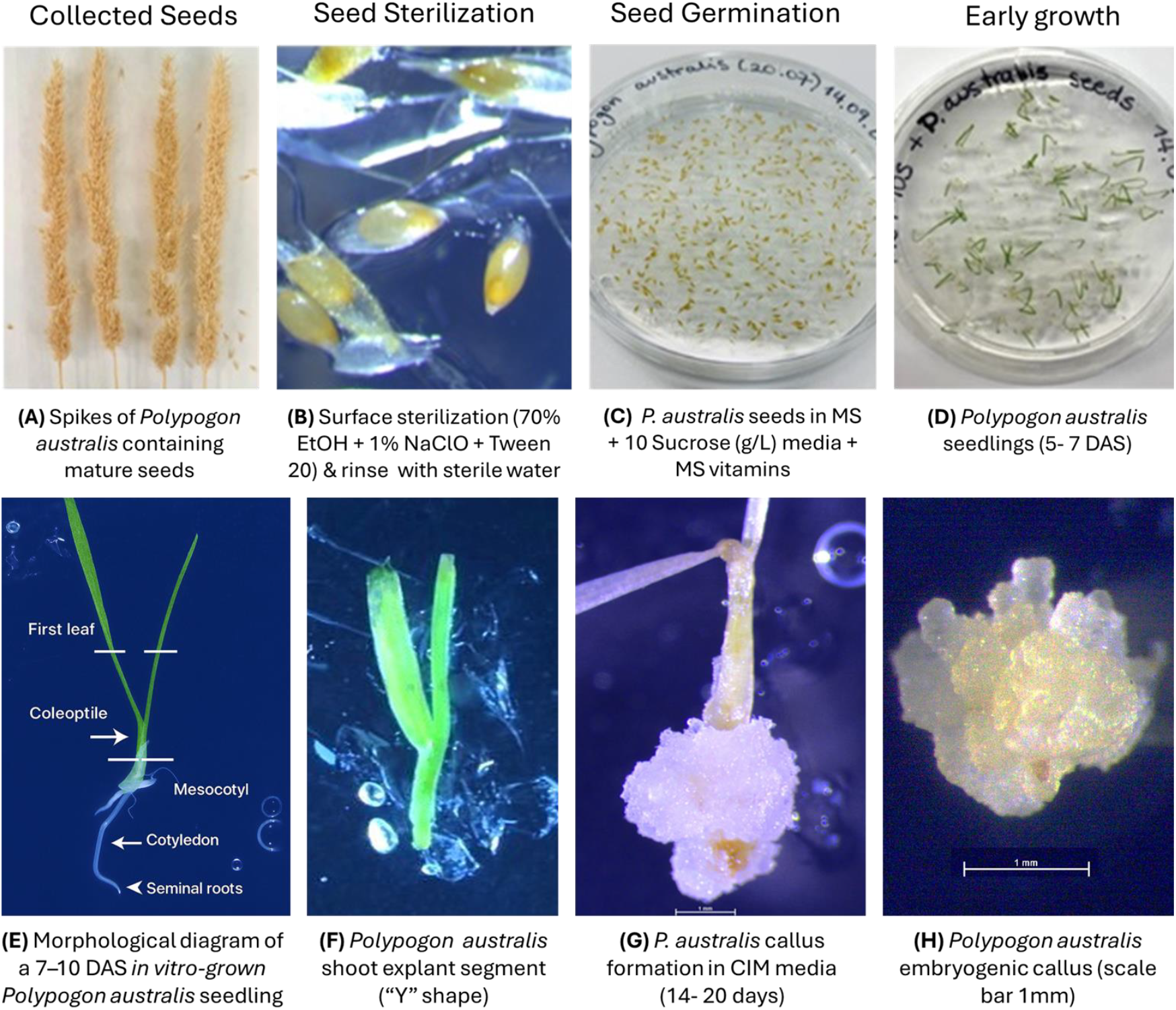
Workflow for *in vitro* seed germination and callus induction in *Polypogon australis*. (A) Spikes collected from field-grown plants; (B) surface sterilization with 70% (v/v) ethanol followed by 1% (w/v) sodium hypochlorite + Tween^®^ 20; (C) seeds cultured on MS medium supplemented with 10 g L^−1^ sucrose and MS vitamins; (D) germination observed within 5–7 days; (E) morphological diagram of a 7–10 DAS *in vitro*-grown seedling highlighting key anatomical structures: first leaf, coleoptile, mesocotyl, cotyledon, and seminal roots; (F) ‘Y’-shaped coleoptile–mesocotyl explants excised from seedlings; (G) explants cultured on callus induction medium (CIM), producing callus tissue within 14–20 days; (H) excised callus (scale bar: 1 mm).

Sterilized seeds were cultured on MS basal medium [18] supplemented with 1% (w/v) sucrose and 0.43% (w/v) Phytagel^®^ (pH 5.75–5.8), and incubated under controlled conditions (22°C, 16/8 h light/dark photoperiod, 136 μmol m^−2^ s^−1^) (Fig. 1C). Germination rate (%) over time was determined in MS+10S medium [19]. Germination percentage was calculated as:

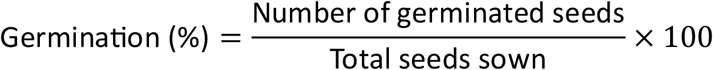

Cumulative germination (%) was recorded at 8, 11, and 15 days after sowing (DAS). The experiment was performed with four independent biological replicates.

### Early growth of P. australis seedlings

Early growth was evaluated by germinating 400–500 mg of surface-sterilized seeds on MS medium (Fig. 1D). After 14 days, five seedlings were transferred into 1-L jars containing MS + 10S medium, and growth was monitored for 50 days. This experiment was conducted with three replicates, each with five plants.

### Callus induction and regeneration efficiency

Coleoptile–mesocotyl junction explants (~0.5 cm segments excised from 7–10-day-old seedlings after emergence of the first true leaf) were cultured on callus induction medium (CIM; MS basal salts supplemented with 2.5 mg L^−1^ Dicamba, 11.3 µM) (Fig.1E). Roots were removed and discarded before culture. Explants were incubated at 24°C in complete darkness. The most responsive explant type was identified after 12 days of culture (Fig. 1F).

Explants were subcultured onto fresh CIM at 3, 4, or 5-week intervals to evaluate callus induction and regeneration efficiency (Fig. 1G-H). Callus induction percentage (CI) was calculated as:

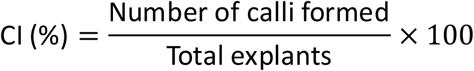

For regeneration, explants were incubated in the dark at 24°C for 2 weeks. Calli were then classified by size, morphology, and color. Embryogenic calli (EC) were excised and subcultured onto CIM for 1 week. Regeneration percentage was calculated as:

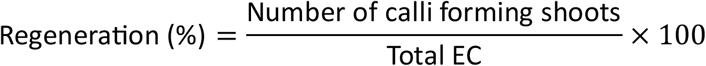

The experiment was performed in triplicate, with 12 explants per replicate.

### Statistical analysis

Germination data were normalized to percentage values (germinated seeds/total seeds × 100) and modeled by fitting a cumulative Gaussian (inverse normal) curve to describe germination dynamics over time. Statistical analyses were performed using GraphPad Prism (version 8.0.1, GraphPad Software, San Diego, CA, USA). A one-way analysis of variance (ANOVA) followed by Tukey’s multiple comparison test was used to determine significant differences among treatments, with p < 0.05 considered statistically significant. Different letters indicate statistical differences between treatments (mean ± SD).

## Results

### Germination and early growth

Germination initiated within 5 DAS on MS medium supplemented with 1% (w/v) sucrose and MS vitamins (Fig. 2A). Approximately 50% of seeds germinated within 16 DAS, with significant differences between 8 DAS and the later timepoints of 11 and 15 DAS. A germination rate of 44–51% is consistent with reports for other native Poaceae that require specific germination conditions, including *Bouteloua curtipendula, Cenchrus ciliaris, Echinochloa crusgalli*, and *Rhynchelytrum repens* [20].

**Figure 2.**
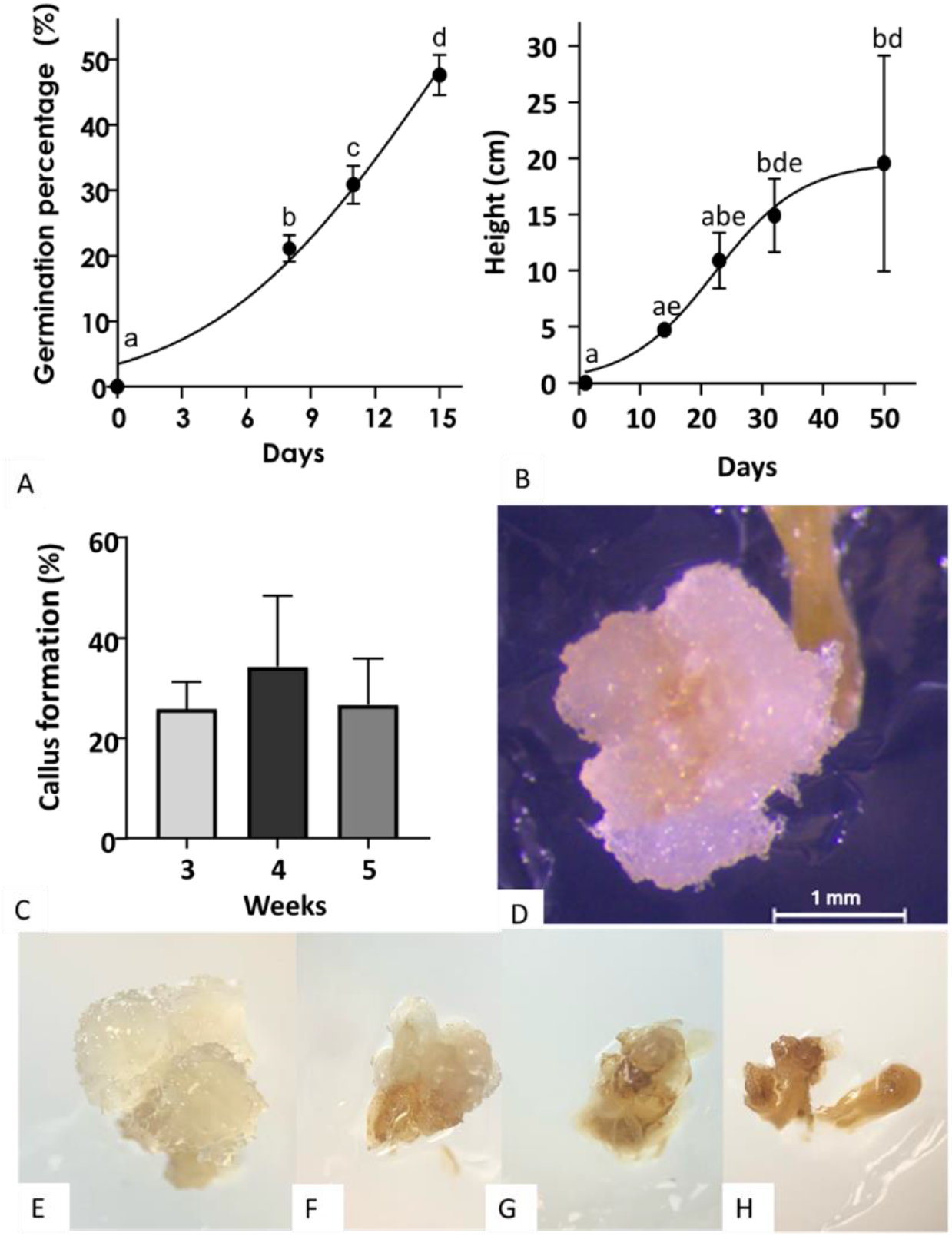
Germination, early growth and callus induction in *Polypogon australis* under controlled conditions. (A) Germination rate (%) over time in MS+10S medium (n=4; Mean ± SD = 47.67 ± 3.15). (B) Seedling height (cm) after germination in MS+10S (n=3). (C) Callus induction (%) from shoot explants cultured in CIM for 3, 4, and 5 weeks (n=5; Mean ± SD = 30.55 ± 11.96). (D–H). Representative images of callus morphology: (D) mesocotyl-derived callus (scale bar = 1 mm); (E, F) friable callus (2 mm and 1.5 mm, respectively); (G, H) compact callus (1 mm and 0.5 mm, respectively). Statistical analyses were performed using one-way ANOVA, and Tukey’s test was applied to determine significant differences (P ≤ 0.05). Different letters indicate statistical significance, while the absence of letters denotes no significant difference.

Seedlings transferred to MS + 10S medium reached an average height of 19.55 cm (SD ± 9.58) at 50 DAS without the need for subculturing (Fig. 2B). Early (1–14 DAS) and late (32– 50 DAS) growth measurements differed significantly, confirming that MS medium supplemented with sucrose effectively supports early development of *P. australis*. These results are consistent with other *in vitro* studies on Chilean Poaceae [16].

### Callus induction and regeneration efficiency

Coleoptile–mesocotyl junction explants were the most responsive tissue for callus induction, with an average induction rate of 30.55% (SD ± 11.96%) after 3–5 weeks in culture (Fig. 2C). Callus size increased with culture duration, reaching 1–3 mm within day 30 (Fig. 2D). Dicamba treatment produced both friable embryogenic calli (EC; 69.33% ± 17.53%) and compact non-embryogenic calli (NEC; 30.67% ± 17.53%) (Fig. 2E–H). Friable calli, characterized by their loose texture, are typically more responsive to somatic embryogenesis, consistent with observations in cereals and forage grasses [21–23].

Comparable behavior has been reported in *Chloris gayana cv*. ‘Tolgar,’ where mesocotyl explants produced embryogenic callus in 3.89% of segments after 5–8 weeks, with four out of eleven induced lines successfully regenerating into plantlets [24]. The use of mesocotyl tissue has been proposed as a strategy to recover totipotent lines and reduce recalcitrance within heterogeneous seed populations, reinforcing its suitability as an explant source. Together, these results support the conclusion that coleoptile–mesocotyl explants are highly effective for embryogenic callus induction in *P. australis*.

### Regeneration potential

Plantlet regeneration was observed 2 weeks after transferring ECs to CIM, showing a regeneration frequency of 45.5% (Fig. 3). Detached ECs produced radicles and cotyledons without the need for exogenous organogenesis-stimulating hormones. Subsequent leaf development occurred within 14 days, and transferring regenerants to MS + 10S medium promoted shoot and root elongation, producing 5–7 cm plantlets by day 30 (Fig. 3). Each EC generated multiple cotyledons, yielding more than one plantlet per explant.

**Figure 3.**
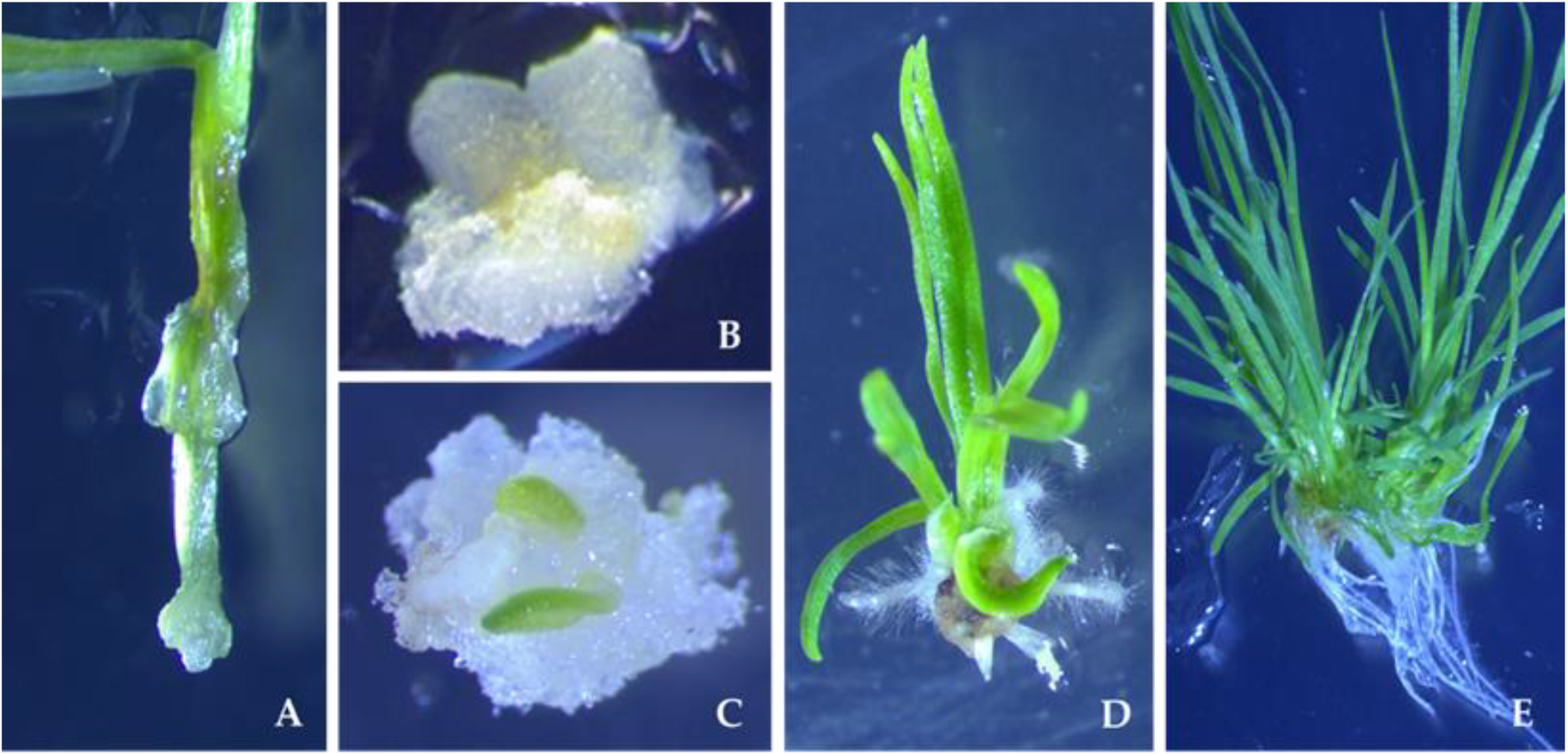
*In vitro* growth and development of *Polypogon australis* plantlets derived from embryogenic calluses. (A) Coleoptile–mesocotyl explant showing early callus formation at 7 days; (B) callus tissue after 10 days of culture; (C) callus with emerging vegetative structures between 14–21 days; (D) detached plantlets regenerated from embryogenic callus cultured on MS medium supplemented with 10 g L^−1^ sucrose, displaying shoot and root formation at 21 days; (E) plantlets after 30 days of growth from the same embryogenic callus.

These results demonstrate the totipotency of *P. australis* ECs and highlight their potential for use in genetic transformation studies. The regeneration efficiency reported here is comparable to or higher than that obtained in other cereal and forage species, where genotype, explant type, and medium composition strongly influence outcomes [25].

## Discussion

This study represents the first report of an efficient *in vitro* regeneration protocol for *Polypogon australis*, a native facultative metallophyte of Chile. The establishment of reproducible conditions for callus induction and regeneration is a crucial step for advancing both basic and applied research in this species. Germination rates (44–51%) and the subsequent development of embryogenic calli from coleoptile–mesocotyl explants are consistent with results obtained in other Poaceae, where this tissue is frequently identified as the most responsive for somatic embryogenesis [21–23]. The 45.5% regeneration efficiency reported here demonstrates the high morphogenic potential of *P. australis* and underscores its suitability for downstream applications such as *Agrobacterium*-mediated transformation.

The results align with studies in related grasses, including *Chloris gayana* and *Echinochloa crusgalli*, where mesocotyl tissues exhibited superior embryogenic responses compared to other explant types [23,24]. Similar protocols have been validated in agronomic and forage grasses, reinforcing the value of coleoptile–mesocotyl explants for overcoming the recalcitrance commonly observed in native species [25,26]. In contrast, regeneration frequencies in other Chilean endemics, such as *Alstroemeria pelegrina* and *Senecio nutans*, remain substantially lower, highlighting the relative efficiency achieved in *P. australis* [27,28].

Notably, the present protocol provides a platform not only for mass propagation but also for functional genomics. Embryogenic calli derived from coleoptile–mesocotyl explants offer a uniform and totipotent tissue type, making them ideal for transformation experiments. This is particularly relevant for CRISPR-based strategies aimed at activating stress-responsive or metal homeostasis genes [7,29]. By integrating these biotechnological tools with the robust *in vitro* system established here, it becomes possible to enhance traits such as Cu uptake, oxidative stress tolerance, and biomass accumulation in *P. australis*, thereby amplifying its potential for phytoextraction or other phytotechnologies.

The environmental context underscores the urgency of this research. Chile remains the world’s largest copper producer, with more than 750 registered abandoned tailings deposits posing risks to soil and water resources [30–32]. Despite mandatory stabilization of mine tailing dams, sustainable long-term remediation strategies remain urgently required [32]. Phytotechnologies utilizing native grasses, such as *P. australis*, offer a cost-effective and environmentally friendly approach [3,4]. Yet, large-scale propagation has been the major bottleneck to their application [5]. By overcoming this barrier, the methodology reported here positions *P. australis* as a model for phytotechnological solutions to address the environmental impacts of mining in Chile and beyond.

## Conclusion and Future Prospects

This work established an efficient and reproducible *in vitro* regeneration protocol for *Polypogon australis*, a native Chilean metallophyte naturally adapted to copper-contaminated soils. Optimized germination and the use of coleoptile–mesocotyl explants enabled high callus induction and regeneration frequencies, overcoming a significant limitation for the large-scale propagation of this native species. This methodology not only supports its application in phytoremediation but also provides a platform for functional genomics and biotechnological approaches—such as CRISPR-dCas9-based activation—to enhance stress tolerance and metal accumulation.

## Ethics approval

Not applicable.

## Consent to participate

Not applicable.

## Consent for publication

Not applicable.

## Availability of data and materials

Not applicable.

## Author contributions

J.V. designed and conducted the experiments, analyzed the data, and wrote the manuscript. C.O. contributed to data analysis and interpretation. G.H. provided technical advice on tissue culture and molecular strategies. R.G. supervised the project and contributed to manuscript revision. All authors read and approved the final manuscript.

